# Incidental identification of maternal malignancies in two Asian women underwent noninvasive prenatal test

**DOI:** 10.1101/095364

**Authors:** Xing Ji, Fang Chen, Yafeng Zhou, Jia Li, Yuying Yuan, Yu Mo, Qiang Liu, Jen-Yu Tseng, Diego Shih-Chieh Lin, Shu-Huei Shen, Yu Liu, Weiping Ye, Hongyun Zhang, Na Liu, Li Shen, Xin Jin, Pi-Lin Sung, Mao Mao

## Abstract

Noninvasive prenatal test (NIPT) has been widely used as a screening test for trisomy 13, 18 and 21 worldwide. Recently, coexistence of maternal malignancy and pregnancy has drawn increasing attention in NIPT studies. Malignancy in pregnant women potentially affected NIPT results, which may cause false positive results or failed tests. However, no such case has ever been reported in Asian population. In this study, for the first time, we reported a stage III dysgerminoma and advanced gastric cancer during pregnancy accidentally identified during NIPT tests. These two women showed aberrant chromosome aneuploidies in NIPT results and concordant pattern of genome disruption found in tumor samples. The findings in this study further validate the effect of maternal malignancy on NIPT results and strengthen the possibility of detecting malignant tumors through NIPT in the future.

## Introduction

Over the past few years, noninvasive prenatal testing (NIPT) has become a common technology screening for the common fetal aneuploidies, including 13, 18 and 21–trisomy. The high sensitivity and specificity of NIPT has been demonstrated by large-scale studies conducted in different populations(Chiuet et al., 2011; Zhang et al.2015). Moreover, NIPT is recommended as a better substitute to traditional Down syndrome screening.

However, despite the high accuracy of NIPT, false or even failed test results still exist in a large amount due to the rapid increasing population who underwent NIPT. The discordance between cell-free DNA (cfDNA) and fetal karyotype could be a major concern of NIPT technic, which could be attributed to various factors including confined placental mosaics, co-twin demise, maternal chromosomal mosaics and maternal malignancy(Bianchi et al., 2015).

Of them, maternal malignancy has drawn more and more attention, due to its fatal consequences during pregnancy. Despite the relatively low incident rate of maternal malignancy (1:1000 to 1: 5000), incidental discovery of maternal cancer have been reported among pregnant women undergoing routine NIPT screening(Amant, Verheecke, et al., 2015; Bianchi et al., 2015; Osborne et al., 2013). However, all of those studies were conducted in western population. Rare case had been reported in Asian population, which is inconsistent with the rapidly increasing number of pregnant women who took NIPT screening. In addition, the majority of maternal malignancies discovered by NIPT are hematological malignancies, few cases had been reported on other cancer types.

In the current report, we presented two cases of aberrant chromosomal aneuploidies unraveled by NIPT subsequently diagnosed with dysgerminoma and gastric cancer, respectively. These case reports supported the potential of applying NIPT to pre-symptomatic detection of malignant tumors in pregnant women in the future.

## Methods and materials

### Sample collection, sequencing and bioinformatics analysis

Ten millimeters of maternal peripheral blood were collected in Cell Free DNA BCT^®^ blood collection tube and was processed within 4 days of collection. Details of the NIPT method, also called Non-invasive fetal trisomy test (NIFTY), have been published previously(Dan et al., 2012). In brief, plasma was separated by sequential centrifugations of the blood sample at 1600g at 4°C for 10mins. Cell free DNA was extracted from plasma and subjected to library construction. The quantity and quality of the library were examined by real time PCR and size distribution. Qualified library was sequenced and the data generated was analyzed using the bioinformatics algorithm to detect fetal chromosomal aneuploidy and large deletion/duplication, as previously described(Chen et al., 2013; Lau et al., 2012).

### Somatic copy-number alterations (Affymetrix OncoScan FFPE Express 2.0)

Somatic Copy number analysis for this tumor was performed using MIP array technology (Affymetrix OncoScan FFPE Express 2.0) with 334,183 sequence tag site probes to measure DNA copy number variations(Wang et al., 2009). Copy number data were processed and normalized by Affymetrix as previously described with passed Affymetrix quality control metrics(Wang et al., 2009). Copy numbers were estimated with the NEXUS software and only samples that passed Affymetrix quality control metrics (median absolute pairwise difference [MAPD] value of□≤□0.6) were considered(Darvishi, 2001).

## Cases summary

### Case 1

The first patient is a 30-year-old woman, who was diagnosed with bilateral ovarian endometriosis and received laparoscopic cystectomy and 6-months gonadotropin-releasing hormone agonist treatment from December 2013 to June 2014. During her first obstetrics ultrasound, bilateral endometriosis, including left ovarian cyst (2.8 cm and 2.0 cm) and one right ovarian cyst (7.6 × 4.4 cm) were diagnosed according to her prior history. The patient received two NIPT tests at 13 and 32 gestational weeks respectively. After the failure of the first test, the second test after resampling of peripheral blood failed again, which excluded the possibility of sample degradation during transportation. To rule out fetal aneuploidies, amniocentesis and array comparative genomic hybridization (array CGH) were conducted on the amniotic fluid, showing a normal fetal karyotype (46, XX). A fetal anatomic ultrasound survey also demonstrated normal fetal anatomy and growth with normal placenta appearing at 19 gestational weeks.

However, the right ovarian cyst with internal flow enlarged gradually as gestational week and fetal MRI revealed that an 8.7×7.9×6.0cm lobulated soft tissue mass at right adnexa with engorged regional collateral vessels noted with diffusion restriction (Figure 1 A and B). At 36 weeks gestational age, patients underwent a cesarean section combined with a conservative debulking surgery, including right salpingo-oophorectomy, right pelvic lymph node sampling, omentectomy and appendectomy. A female infant with Apgar score 5/8 and weight of 2498 g appeared normal without any obvious dysmorphic features. The final pathologic diagnosis was a right ovarian dysgerminoma, pT3aN0M0, FIGO IIIa with tumor invasion to appendix (Figure 1C and D). Thereafter, the patient received 6-course of chemotherapy (bleomycin, etoposide and cisplatin).

**Figure 1.**
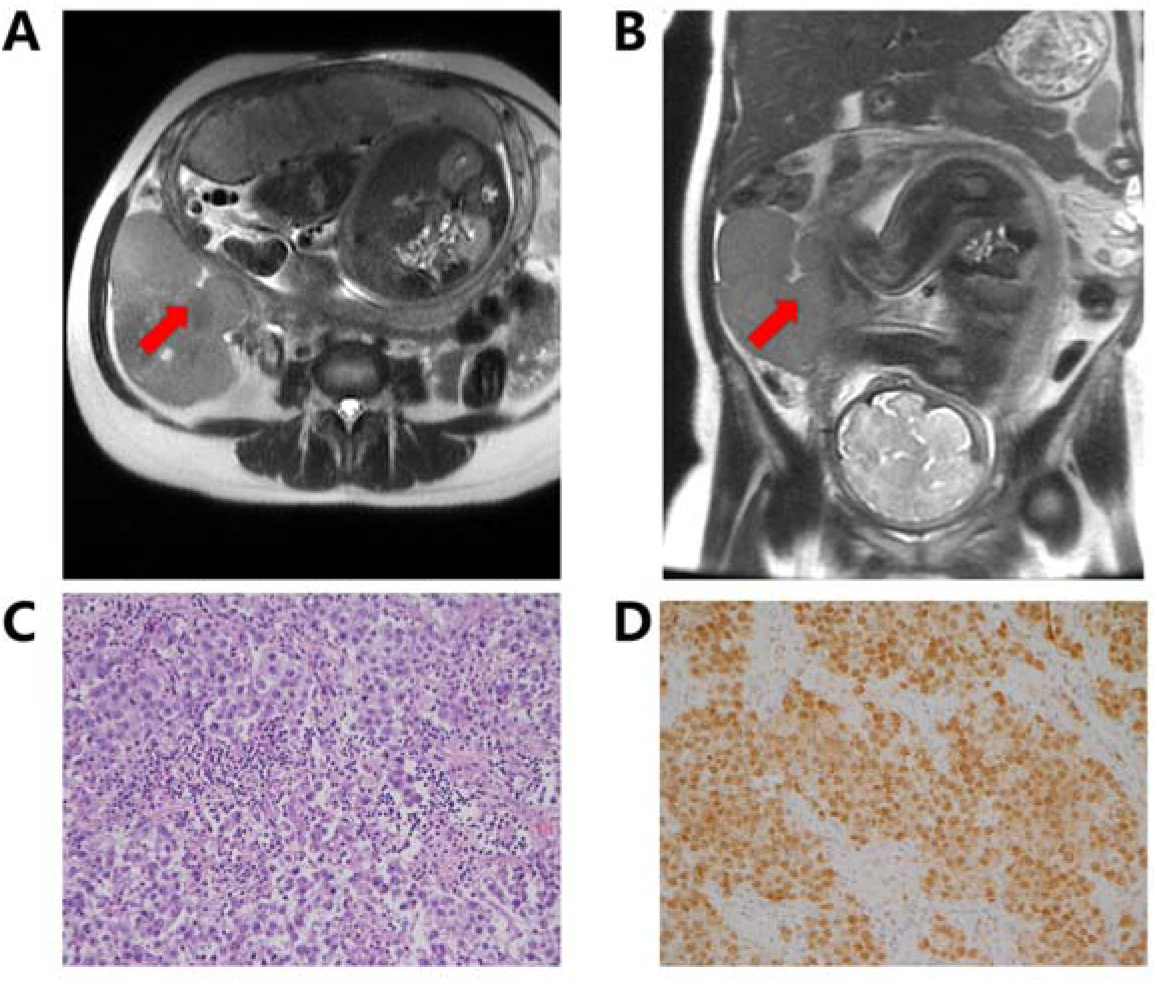
Dysgerminoma in a 31-year-old pregnant female patient. A 112×109×75 mm mass on T2-weighted MRI images at 35 gestational weeks (A: axial plane, B: coronal plane). The red arrow heads indicate the malignant tumors in all panels. (C) solid sheets of large tumor cells with prominent nucleoli and clear cytoplasm (H&E staining, x 200). (D) the tumor cells were immuno-reactive for OCT4 (anti-OCT4, x200).

After knowing concurrent pregnancy and dysgerminoma in this patient, we re-examined the NIPT results and found multiple aneuploidies in the blood DNA, which was the main reason of NIPT failure (Figure 2A).

**Figure 2.**
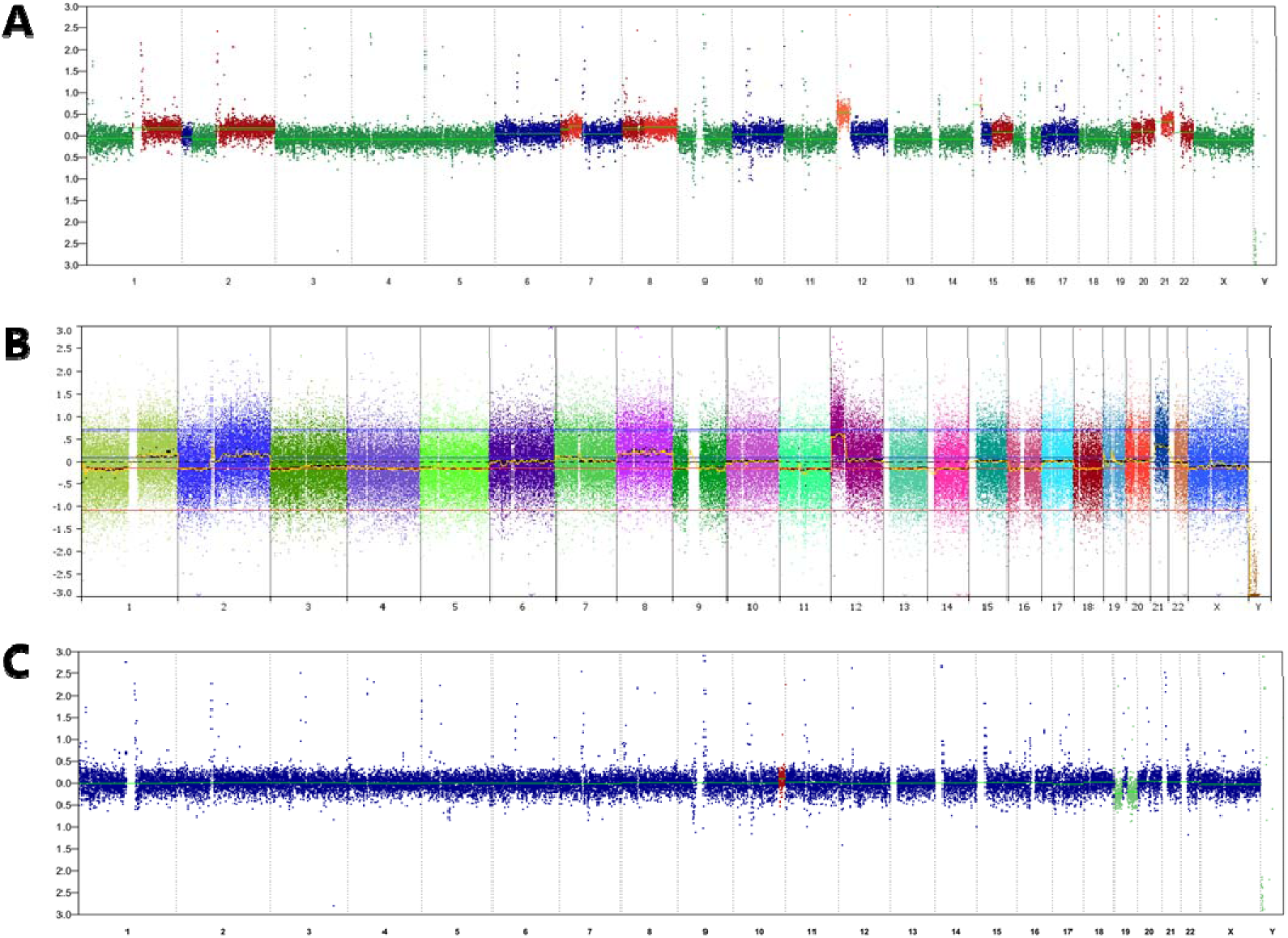
Whole-genome view of copy-number gains and losses in plasma (A), malignant tumors (B) and plasma samples after treatment (C).

To study the resource of genetic variations found in NIPT, MIP array technology (Affymetrix OncoScan FFPE Express 2.0) with paraffin-embedded tumor tissues was performed. We found the concordance of chromosome alterations between maternal plasma DNA and tumor tissue (Figure 2A and B), such as gain of whole short arm of chromosomal 12 (12p11.1-13.3) and gain of whole chromosomal 21 (p11.2–q22.3). Those results suggested that tumor cfDNA contributed to the aberrant NIPT results.

To further validate this result, another cfDNA test was conducted after the surgery and six courses of chemotherapy. As shown in figure 2C, aberrant copy number variations disappeared after the complete treatment, further supporting the hypothesis that NIPT could detect tumor cfDNA in maternal plasma. These results strengthened the potential implement of non-invasive prenatal test in screening and monitoring maternal cancer during pregnancy.

### Case 2

The second patient is a 36-year-old woman, who received two NIPT tests at 16 and 20 gestational weeks, respectively. The patient had peptic ulcer history, she suffered from growing nausea and vomiting during the pregnancy. The patient received a gastroscope and was diagnosed with an advanced gastric cancer (Figure 3). After one week of diagnosis, she died of gastric cancer. Due to lack of cancer samples, no experiment was performed to validate the copy number changes in gastric cancer tissues. A NIPT test at 30-fold depth, which equals to 3X WGS, was performed using the NIPT DNA library. Both original NIPT and the 30-fold NIPT results displayed a severe disruption of genome stability, further supporting abnormal copy number variations reported by NIPT test is a strong indicator of maternal malignancies (Figure 4). In line with previous reports (CALCAGNO et al., 2005; Network, 2014), genomic changes commonly seen in gastric cancer, such as aneuploidy of chromosome 7 and 8, were observed, providing further evidence that NIPT test is able to detect the tumor ctDNA in maternal blood.

**Figure 3.**
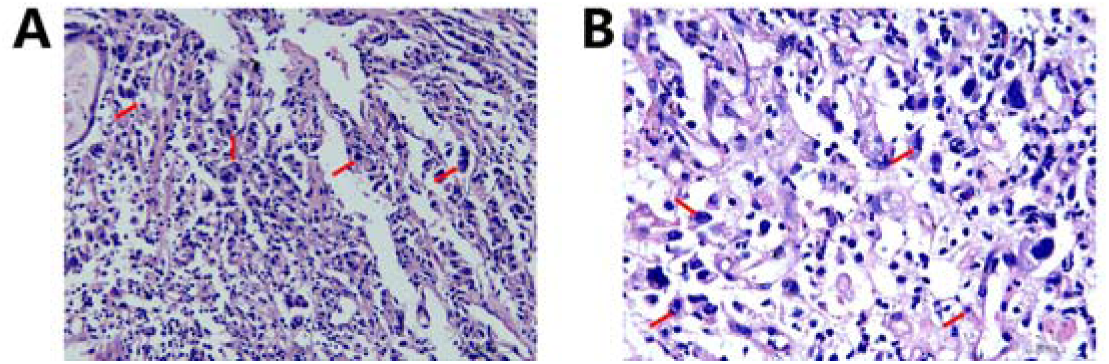
A gastric cancer in patient 2. The red arrowhead marks signet ring cells in gastric cancer biopsy samples (A: H&E staining, x 20, B: H&E staining, x 40).

**Figure 4.**
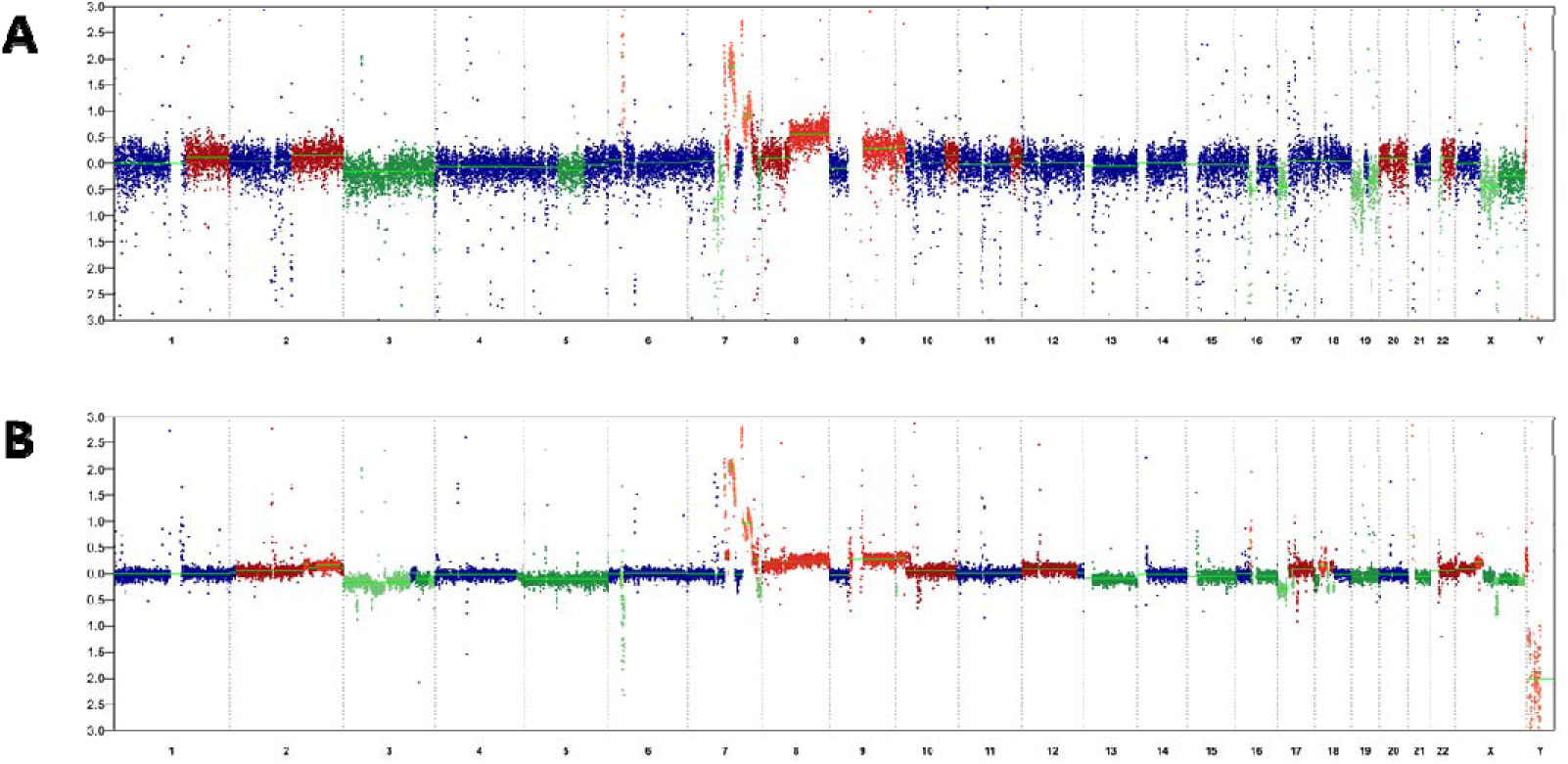
Whole-genome view of copy-number alterations in plasma revealed by NIPT (A) and 30-fold NIPT (B).

## Discussion

Incidental discovery of maternal cancer has been repeatedly reported among pregnant women undergoing routine NIPT screening in Europe and US (Amant, Vandenbroucke, et al., 2015; Bianchi et al., 2015; Osborne et al., 2013). Although different groups have reported their findings of maternal cancer in pregnant women undergoing NIPT screening, our report is the first to reveal the association between maternal cancer and aberrant NIPT results in Asian population. In this study, two Asian pregnant women showed aberrant NIPT results and subsequently diagnosed with maternal malignancies, both harboring extensive copy-number changes across the whole genome. According to previous study(Bianchi et al., 2015), patients with multiple aneuploidies are strongly associated to maternal cancer, which is also confirmed by our results.

Moreover, further investigation of NIPT result, in combination of SNP array outcome demonstrated similar copy-number variation patterns in samples from peripheral blood and tumor tissue, which provides strong evidence for the detection of circulating cancer DNA from blood using extreme-low depth whole genome sequencing.

The most frequently diagnosed malignancies during gestation are breast cancer, cervical cancer, Hodgkin’s disease, malignant melanoma, and leukemia (Pavlidis, 2002). Previous reported maternal malignancies contain a large proportion of blood cancer(Amant, Verheecke, et al., 2015; Bianchi et al., 2015). For the first time, we showed a stage III dysgerminoma and advanced gastric cancer during pregnancy, further expanding the range of cancer types detected by NIPT screening.

According to previous reports, the incidence rate of cancer complicating pregnancy ranges from 1:1000 to 1: 5000 among pregnancies (Pavlidis NA, 2002) whereas the reported incidence of maternal cancer associated with abnormal NIPT results is much lower, which is the case in our study. So far, only two cases of abnormal NIPT results were confirmed with concurrent maternal malignancies, despite the large number of NIPT tests conducted in Asia. One possible explanation is that present NIPT products focus mainly on the abnormal trisomy-13, 18, and 21, multiple aneuploidies were shown to be frequently correlated with maternal cancer whereas in NIPT screening, it often causes screening failure thus may be overlooked. Another possibility is that maternal malignancy is a relatively rare disease often misdiagnosed by other pregnant symptoms and results in a low diagnostic rate. The complicated conditions during pregnancy makes it even more difficult to perform medical follow-up for patients. Many patients may have concurrent cancer during pregnancy after their NIPT tests and do not report to their NIPT providers. Therefore, more attention and medical follow-up are greatly needed for those patients who have multiple aneuploidies in NIPT test. However, how to conduct reasonable clinical follow-up and provide proper medical interventions becomes an important question that should be answered by NIPT provider and medical staff in the future.

Of these two patients described herein, one patient gave birth to a healthy girl and subsequently underwent successful resection of tumor tissue, followed by chemotherapy. The other patient died one week after diagnosis with gastric cancer. The limitation of this study is that we failed to obtain enough tissue samples for the second patient to discover tumor DNA information.

Taken together, our findings underlined potential value of NIPT for pre-symptomatic cancer screening in pregnant women, indicating a possible direction of early cancer screening strategy.

